# The dynamic response of quorum-sensing to density is robust to signal supplementation and signal synthase knockouts

**DOI:** 10.1101/2022.09.12.507654

**Authors:** Jennifer B. Rattray, Patrick Kramer, James Gurney, Stephen Thomas, Sam P. Brown

## Abstract

Quorum sensing (QS) is a widespread mechanism of environment sensing and behavioral coordination in bacteria. At its core, QS is based on the production, sensing and response to small signaling molecules. Previous work with *Pseudomonas aeruginosa* shows that QS can be used to achieve *quantitative* resolution and deliver a dosed response to the bacteria’s density environment, implying a sophisticated mechanism of control. To shed light on how the mechanistic signal components contribute to graded responses to density, we assess the impact of genetic (AHL signal synthase deletion) and/or signal supplementation (exogenous AHL and PQS addition) perturbations on *lasB* reaction-norms to changes in density. Our approach condenses data from 2,000 timeseries (over 74,000 individual observations) into a comprehensive view of QS-controlled gene expression across variation in genetic, environmental, and signal determinants of *lasB* expression. We first confirm that deleting either (Δ*lasI*, Δ*rhlI*) or both (Δ*lasIrhlI*) signal synthase genes attenuates QS response. In the Δ*rhlI* background we show persistent yet attenuated density-dependent *lasB* expression due to native 3-oxo-C12 signaling. We then test if density-*independent* quantities of signal (3-oxo-C12, C4, PQS or combined) added to the WT either flatten or increase the reaction norm and find that the WT response is robust to all tested concentrations of signal, alone or in combination. We then move to progressively supplementing the genetic knockouts and find that cognate signal supplementation (Δ*lasI*+3-oxo-C12, Δ*rhlI*+C4) is sufficient to restore *lasB* expression and as well as reactivity to density encoded by the intact signal synthase. We also find that dual supplementation of the double synthase knockout restores expression but does not flatten the reaction norm. Despite adding a density-*independent* amount of AHL, the double signal synthase can still quantitively sense density. Our results show that a positive reaction norm to density is robust to multiple combinations of gene deletion and density-independent signal supplementation and that while density-independent signal supplementation can increase mean expression, the WT QS still retains the ability to quantitatively respond to density. Our work develops a modular approach to query the robustness and mechanistic bases of the central environmental *sensing* phenotype of quorum sensing.

## Introduction

Many species of bacteria use a form of cell-cell communication known as quorum sensing (QS) to collectively sense and respond to variation in their extracellular environment. QS bacteria secrete and respond to diffusible signal molecules that encode information on aspects of their environment. Canonically, QS is understood as a density sensing device, as higher density populations will typically accumulate higher concentrations of QS signals, although other aspects of environmental variation can also impact signal supply (e.g. mass-transfer (1–3), and genetic similarity (4)).

In the context of sensing density, the adoption of a “quorum” analogy (5) leads to a simple threshold, qualitative interpretation of QS behavior as either ‘quorate’ (high density; QS-controlled genes ‘on’) or ‘sub-quorate’ (low density; QS-controlled genes ‘off’). However, a growing number of studies have revealed substantial heterogeneity in responses to QS signals (1,6–17) and we recently demonstrated that QS in *Pseudomonas aeruginosa* does not necessarily function in a threshold manner, on both population and single-cell scales ((18). Specifically, we found that per capita *lasB, pqsA*, and *rhlI* expression shows a linear, quantitatively graded expression control (or ‘reaction norm’, (19,20)) on the population scale to variation in density.

The ability of *P. aeruginosa* to achieve quantitative resolution and deliver a dosed response to its social environment implies a sophisticated mechanism of control. QS in *P. aeruginosa* represents one of the most intensely studied model systems – revealing a complex intracellular regulatory network driven by multiple signal molecules (21,22) (Figure 1). Figure 1 illustrates the coupling between signal synthesis, signal response, and downstream expression. The two primary signal molecules of *P. aeruginosa* are the *N*-acyl homoserine lactones (AHLs) N-(3-oxododecanoyl)-L-homoserine lactone (3-oxo-C12-HSL, henceforth 3-oxo-C12) and N-butyryl-homoserine lactone (C4-HSL, henceforth C4). The AHL system in PA is conventionally understood as a hierarchical system (23–27), with the *lasIR* system (producing/responding to 3-oxo-C12) governing the *rhlIR* system (C4), although alternative *las-rhl* wirings have been reported (28,29). On-going discoveries of additional extracellular signals (31) and intracellular regulatory feedbacks are continually revising the mechanistic picture for the highly studied reference strain PAO1 (32). In this study, we focus on *lasB*, a secreted protease and virulence factor that is under dual AHL signal control (Figure 1, (24,33–35). We focus on *lasB* because of its widespread use as a model of *P. aeruginosa* virulence (36,37), cooperation (38–41) and as a marker of QS-controlled behaviors (42–44).

**Figure 1.**
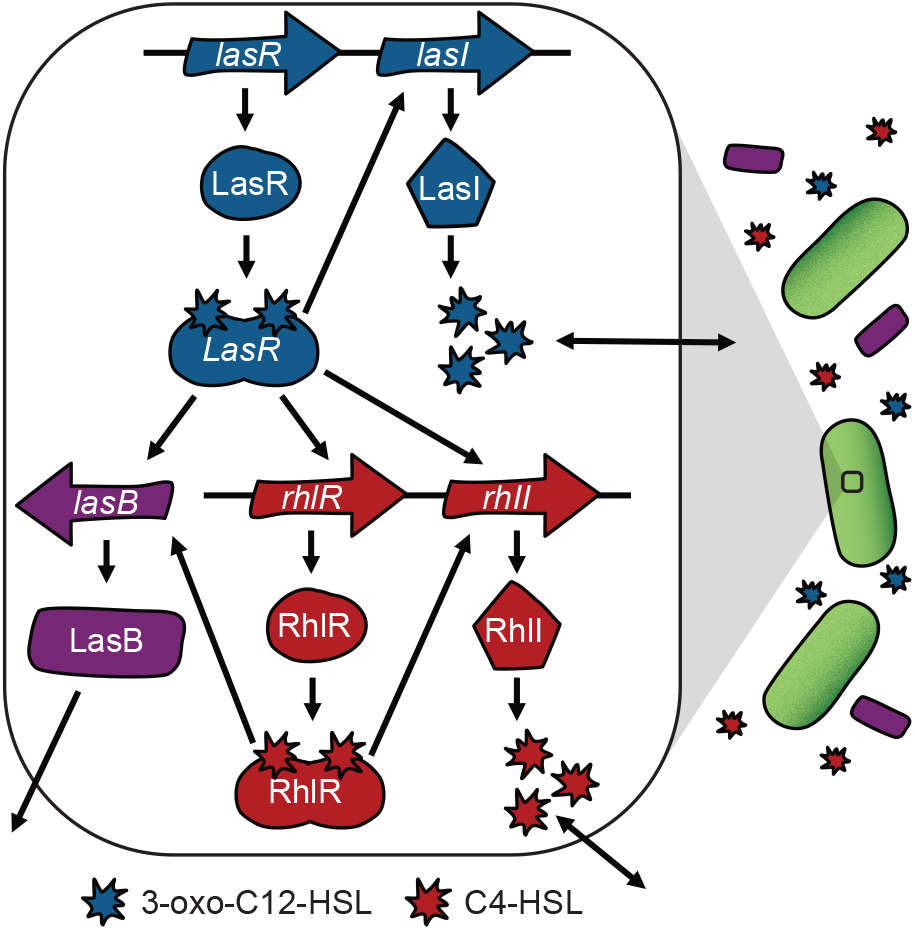
Intracellular mechanics of AHL signaling in *Pseudomonas aeruginosa*. The *P. aeruginosa* QS system is dominated by the las (blue) and rhl (red) acyl-homoserone lactone (AHL) signaling systems. Each system codes for a signal synthase gene (*lasI, rhlI*), which guide the production of a diffusible AHL signal molecule (N-(3-oxododecanoyl)-L-homoserine lactone (3-oxo-C12-HSL, henceforth 3-oxo-C12) and N-butyryl-homoserine lactone (C4-HSL, henceforth C4)) at an initially basal level. Binding of each signal to its cognate receptor (LasR-3-oxo-C12, RhlR-C4) results in an active transcriptional factor which up-regulates cognate synthase activity (signal auto-induction – a positive feedback control of signal production) along with other genes in the QS regulon, e.g. the secreted exoprotease enzyme and virulence factor LasB (purple).

To shed light on how these mechanistic components contribute to graded responses to density, we assess the impact of genetic (AHL signal synthase deletion) and/or chemical (exogenous signal supplementation) perturbations on *lasB* reaction-norms to changes in density. Figure 2 illustrates how our approach differs from conventional knockout/complementation approaches, which focus on the impact on gene expression in a single environmental condition (typically, growth in complex medium to high density; Figure 2.A). In the context of *lasB* expression, studies have demonstrated that *lasB* expression can be attenuated by knockouts of one or both AHL signal synthase gene (*lasI* and/or *rhlI*, Figure 2.A grey circle) and can then be restored by addition of missing signal (Figure 2.A green circle) (45).

**Figure 2.**
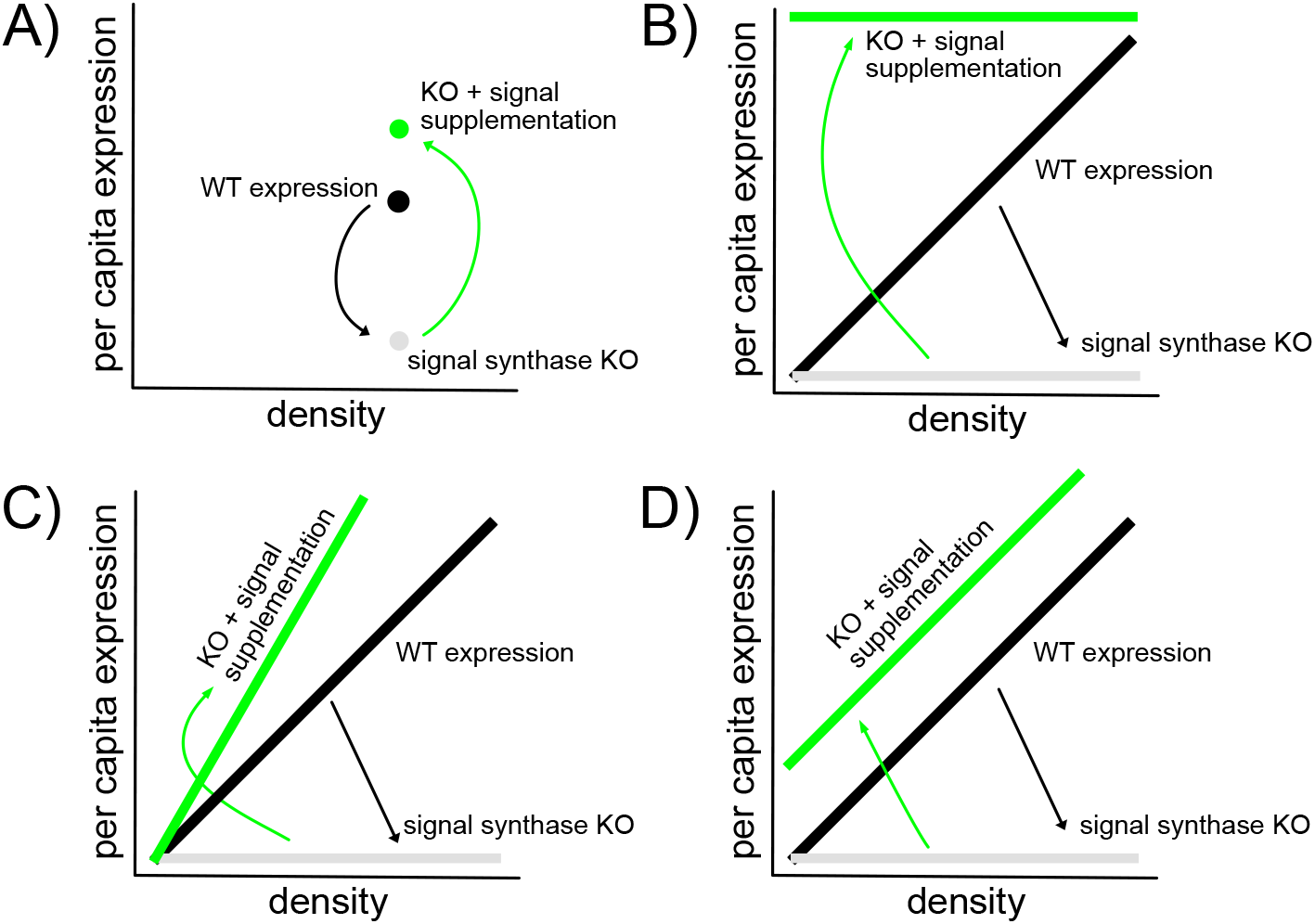
Gene knockout and complementation experiments as a function of environmental variation. A reaction-norm approach (Woltereck 1910; Waddington 1942; Schlichting and Pigliucci 1998; Paaby and Testa 2018; Rattray et al. 2022) introduces more metrics for the impacts of knockout / complementation experiments, including changes in reaction norm slope and average expression level across densities. Black = wildtype per-capita gene expression. Grey = attenuated per-capita gene expression in a signal synthase knockout. Green = genetic knockout supplemented with exogenous signal. A) conventional knockout / complementation experiments are conducted under a single controlled lab condition (for QS experiments, typically high-density environments). B,C,D) Reaction norm experiments conducted under a range of environmental conditions (e.g. differing stationary phase densities). The grey lines represent the hypothesis that knocking out one or both signal synthase gene will ‘flatten off’ the wildtype *lasB* reaction norm shown in the black bars, due to the dependency of *lasB* expression on both AHL signal inputs (Figure 1). B) The “flatten up” hypothesis predicts that sufficient signal supplementation will remove encoding of density information in the signal environment, producing high mean expression and a slope of zero. C) The “acceleration” hypotheses predicts that signal supplementation can steepen the reaction norm compared to the wildtype due to synergistic *lasB* expression control, particularly if one or both native synthases are intact. D) The “robust reactivity” hypothesis predicts that signal supplementation can raise mean expression but retain reactivity (positive slope). In the results section we develop more specific hypotheses leading to predictions of patterns (B,C,D) under defined synthase knock-out and signal supplementation manipulations.

In contrast to the conventional complementation approach focused on expression in a single environment (Figure 2.A), we turn to an examination of how signal knockouts and complementation modify reaction norms, i.e. the *reactivity* of QS controlled *lasB* to changes in the social environment (Figure 2.B,C,D). In light of the synergistic dual signal control model of *lasB* (Figure 1; (24,33,35,46)), we predict that genetic knockouts of *lasI* or *rhlI* alone or in combination will attenuate the QS response, specifically flattening the reaction norm towards an ‘always off’ phenotype (Figure 2.B,C,D, grey lines). Turning to supplementation, the hypothesis in analogy to Figure 2.A is that complementing the signal synthase knockouts (or indeed the wildtype) with sufficiently high concentrations of both signals will lead to an always on phenotype, represented by high *lasB* expression across all density conditions where the slope “flattens-up” (Figure 2.B, green bar).

In light of the synergistic response of *lasB* expression to a dual input of the two AHL signals, we offer an alternate hypothesis that signal supplementation can in some circumstances lead to enhanced reactivity of the wildtype to density (steeper reaction norms, Figure 2.C green bar), due to the amplifying effect of the synergistic *lasB* signal response. Specifically, we predict steeper reaction norms under a combination of a single intact signal synthase (providing information on density) with exogenous supplementation of the knocked-out signal synthase (magnifying the response to density due to synergistic *lasB* expression control). Figure 2.D outlines a final potential pattern, outlining that signal supplementation could increase mean expression, while leaving reactivity (positive slope) intact. This “robust reactivity” hypothesis would indicate that while *lasB* expression is sensitive to exogenous signal, the ability of QS to quantitatively tune responses to changes in density is robust to exogenous signal.

Consistent with the ‘robust reactivity’ hypothesis (Figure 2.D), our results show that a positive reaction norm to density is robust to multiple combinations of gene deletion and density-independent signal supplementation, with little support for the ‘flatten up’ or ‘acceleration’ hypotheses (Figures 2.B,C). We find that density-independent signal supplementation can increase mean expression, but that WT QS still retains the ability to quantitatively respond to density. Finally, a positive reaction norm to density persists even in the absence of *both* AHL synthase genes, given density-independent signal supplementation. We discuss these results in light of the complex regulatory control of *lasB* and in the context of evolutionary theory of communication.

## Results

To provide a baseline for subsequent analyses of genetic knockout and signal supplementation manipulations, we begin with a reaction-norm analysis of NPAO1 wildtype *lasB* expression across a range of population carrying capacities, using an automated analysis approach that extracts growth and expression data from timeseries experiments (Figure 3). Our results recapitulate earlier findings that *lasB* obeys a linear, graded reaction-norm on the population scale (Figure 3.C, (18)) and outlines the high throughput methodological approach we take throughout the rest of this study (Figure 3.A-C), which allows us to assess reaction norms across a large number of genetic and chemical manipulations.

**Figure 3.**
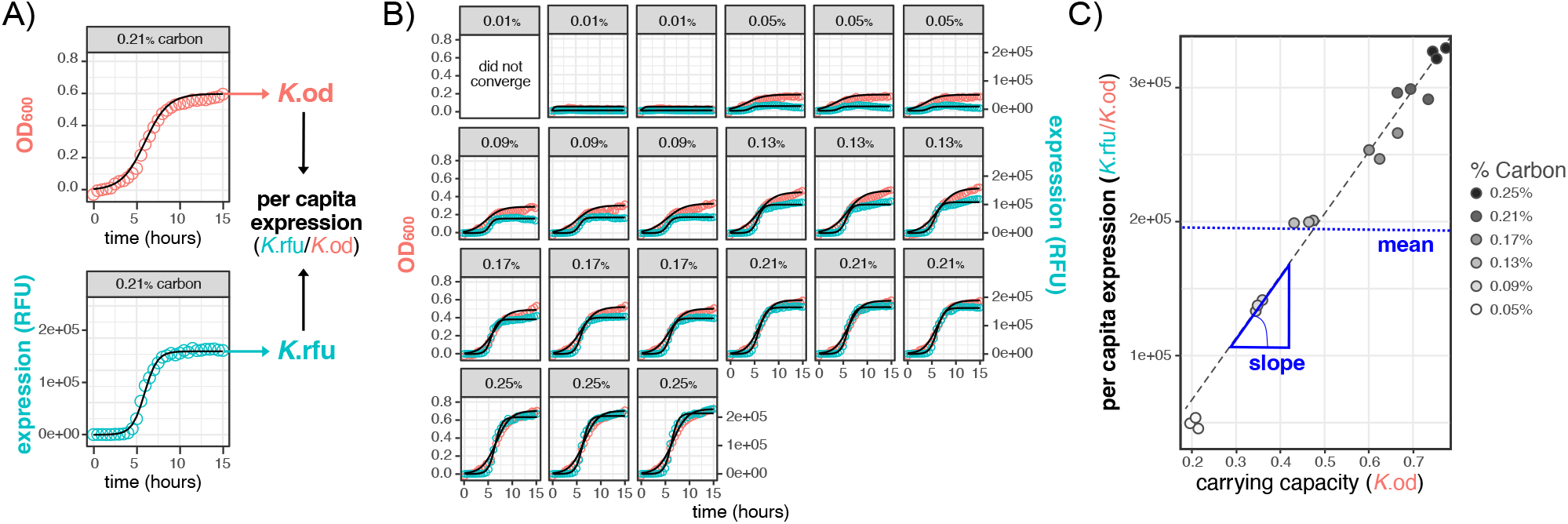
A reaction norm methodology to quantify QS phenotypic responses. In order to dissect broad trends in quorum sensing behavior across population densities, we reduced the dimensionality of our data by first summarizing the growth and *lasB* gene expression of each population (A) across controlled densities (B) and then summarize the data across environmental density (C). A) Logistic curves were fit to raw expression (RFU) and growth (OD600) data of wildtype NPAO1 containing the short half-life *PlasB::gfp*(ASV) quorum sensing reporter (pMHLAS). Cells were grown in triplicate for 20 hours (15 hours shown above for brevity). We then extract the K, commonly referred to as the carrying capacity, from the logistic equation and use a ratio of K_rfu_/K_od_ to describe the per capita behavior of that population. B) Seven distinct culture carrying capacities were generated by manipulating the concentration of casein digest as the limiting resource, the lowest was removed from analysis due to lack of convergence in the logistic fits. C) Per capita expression (K_rfu_/K_od_) is then plotted against carrying capacity (K_od_) and a linear regression (grey dotted line) is performed to generate the reaction norm, i.e., the change in behavior across an environment. This reaction norm can then be further summarized by calculating the slope and mean across density.

### Deletion of signal synthase genes attenuates quorum sensing activity and responsiveness

In our first QS manipulation, we assess the impact of signal synthase genetic knockouts on *lasB* reaction norms. In light of the dual signal control of *lasB* (Figure 1, (23–26,33,35,46), we hypothesized that knocking out any combination of *lasI* and *rhlI* signal synthases will attenuate QS reaction norms, with greater attenuation for *lasI* alone compared to *rhlI* alone due to the hierarchical arrangement of *las* and *rhl* systems (27,28,33,47).

Figure 4 illustrates support for our “flattening off” hypotheses (grey lines, Figures 2B-D). Specifically, we test for decreases in mean expression and reaction norm slope compared to the WT and find that knocking out any gene alone or in combination attenuates both mean QS activity and responsiveness to density (ANOVA, F(3,494) = 1910, p < 2.2e-16. Post-hoc comparisons using Dunnett’s Test (to control for multiple testing), df = (2, 494), one tailed p < 0.001 for all comparisons to WT). Additionally, compared to the double signal synthase knockout, knocking out *lasI* alone reduces QS response to a similar extent as knocking out *lasI* and *rhlI* together (Dunnett’s Test, df = (1, 372), one-tail p_mean_ = 0.4726, p_slope_= 0.6965). In the case of the *rhlI* knockout compared to the double knockout, we still see a residual quorum-sensing response (Dunnett’s Test, df = (1, 372), one-tail p_mean_ < 0.0001, p_slope_ < 0.0001), consistent with partial and density-dependent *lasB* expression in response to natively produced 3-oxo-C12 alone (42).

**Figure 4.**
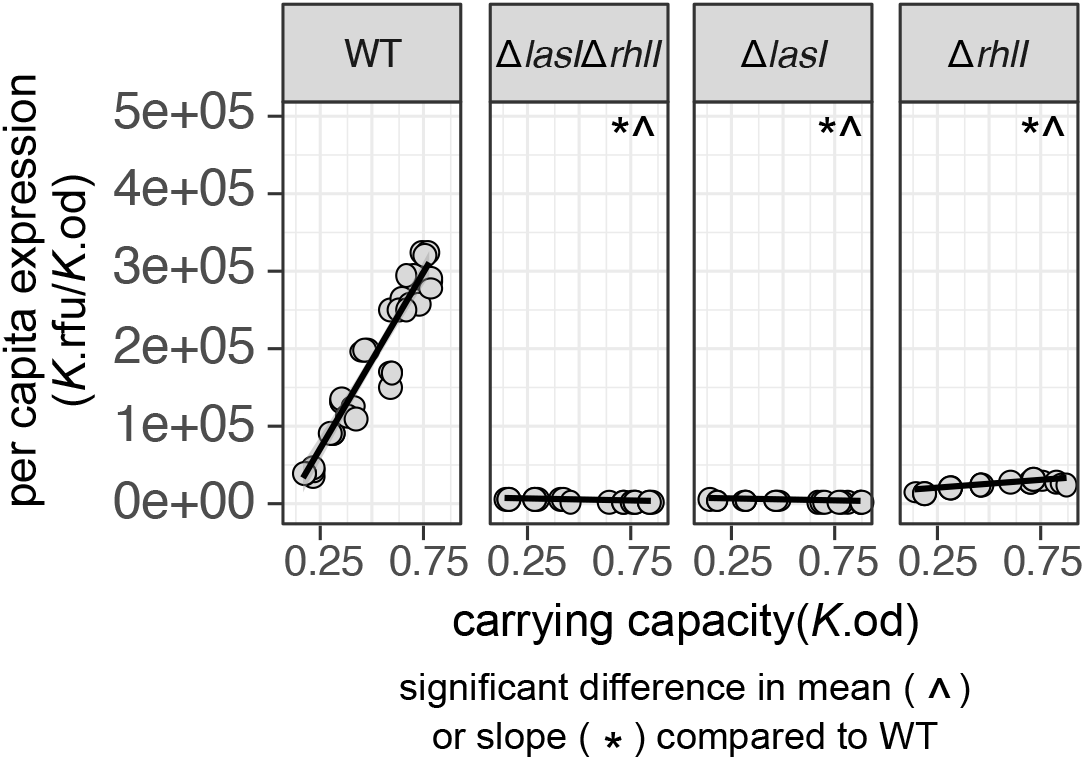
Deletion of signal synthase genes attenuates quorum sensing activity. Reaction norms of per capita *lasB* expression (y-axis) across density (x-axis) in dual whole gene deletion (Δ*lasI*Δ*rhlI*, 3-oxo-C12 and C4 signal synthase knockout) and single whole gene deletion (Δ*lasI*, 3-oxo-C12 signal synthase knockout; Δ*rhlI*, C4 signal synthase knockout) backgrounds. There is a significant reduction in mean expression and slope for all signal synthase knockouts compared to the WT (ANOVA, F(3,494) = 1910, p < 2.2e-16). Post-hoc comparisons using Dunnett’s Test (to control for multiple comparisons), df = (2, 494), one tailed p < 0.001). ^ Indicates a significant decrease in mean compared to the WT. * Indicates a significant decrease in slope compared to the WT.

### Exogenous signal supplementation of the wildtype increases mean response and maintains reactivity to density

We next turn to manipulations to increase signal exposure via exogenous supplementation of 3-oxo-C12 HSL and/or C4 HSL in the wildtype. The AHL signaling molecules used in this study are suspended in methanol, so methanol was used as a control and the amount of methanol used did not significantly impact growth or average expression levels (Supplemental Figure 2). In Figure 5, we find that in most cases adding any fixed (density-*independent*) concentration of either signal, alone or in combination, significantly increases the mean level of expression across the reaction norm compared to the WT with no supplementation (ANOVA, F(15,391) = 106, p < 2.2e-16. Dunnett’s Test, df = (15, 391), one tail p <.0001). The only two exceptions we see are with 3-oxo-C12 (10uM and 50uM of 3-oxo-C12), and in the latter case we see a significant *decrease* of mean expression (Dunnett’s Test, df = (15, 391), two tail p <.0001). This observed decrease could be a result of a high concentration of 3-oxo-C12 (48), which aggregates into detrimental micelles at high concentrations (49) while C4 does not.

**Figure 5.**
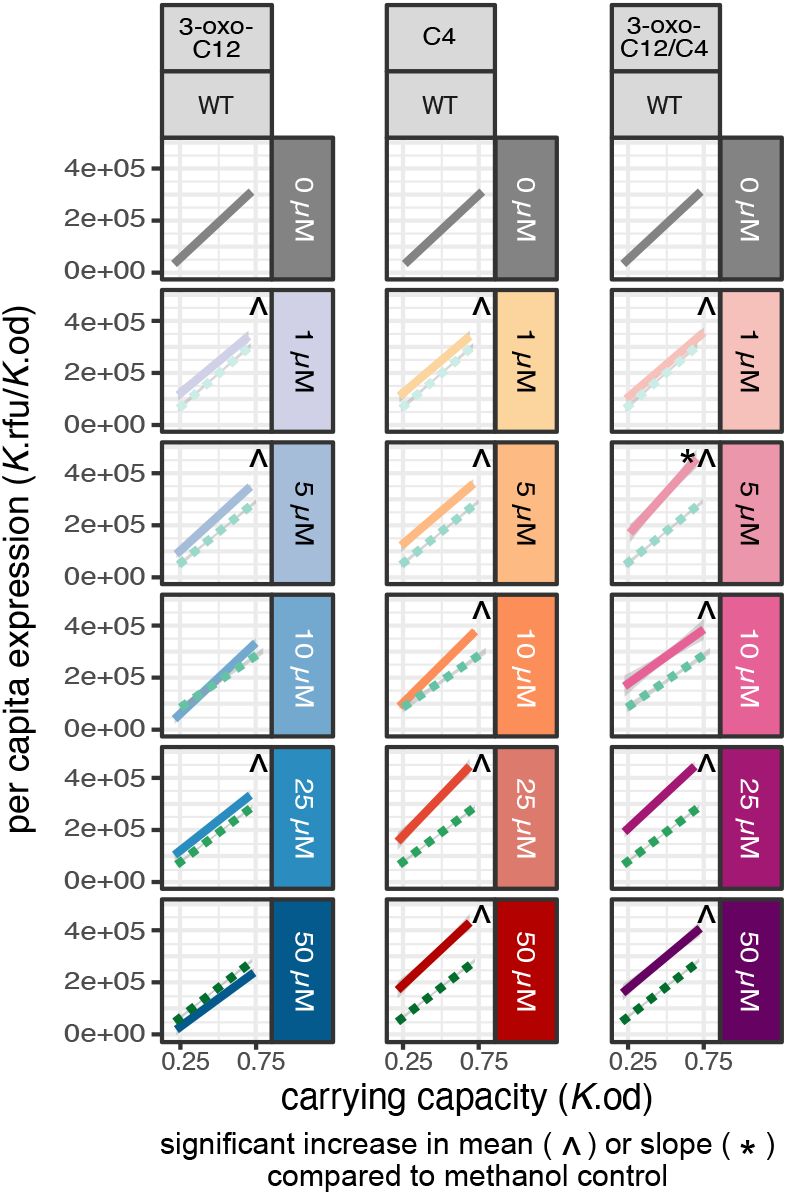
Exogenous signal supplementation of the wildtype increases mean response and maintains reactivity to density. Reaction norms of per capita *lasB* expression (y-axis) across density (x-axis) in the WT quorum sensing background. WT behavior with no signal supplementation in grey. Methanol control (green, dotted line) and signal supplemented environments [3-oxo-C12 alone (blue), C4 alone (orange), 3-oxo-C12 and C4 combined (pink)] across 5 different concentrations of signal (1uM, 5uM, 10uM, 25uM, 50uM). Each reaction norm is built using a linear regression on 18 data points (6 carrying capacity environments run in triplicate). ^ Indicates a significant increase in mean compared to the non-supplemented methanol control. * Indicates a significant increase in slope compared to the non-supplemented methanol control. We find a significant increase in means for all manipulations compared to the non-supplemented WT (ANOVA, F(15,391) = 106, p < 2.2e-16. Dunnett’s Test, df = (15, 391), one tail p <.0001), but only one significant increase in slope (Dunnett’s Test, df = (15, 391), one tail p_5uM-3-oxo-C12/C4_ 0.0023). For legibility, Figure 5 only shows the fitted linear models and not the underlying 18 datapoints per linear fit. See Supplemental Figures 1–2 for a series of plots of the experimental and control data plus linear model fits.

The increase in mean expression in the WT upon signal supplementation indicates that native signal production is not sufficient to maximize gene expression, averaging across densities. But does the addition of fixed amounts of signal modulate the *responsiveness* of the WT to different densities? The hypothetical models in Figure 2 are motivated by gene knockout (grey) plus signal complementation (green) but can be translated to the case of wildtype supplementation. Figure 2.B (green line) represents the hypothesis that exogenous supplementation of (sufficient) signal will maximize QS-controlled gene expression (‘flattening up’), regardless of density, while Figures 2.C,D (green line) represent the hypotheses that augmenting the availability of either signal past WT levels will enhance (Figure 2.C) or maintain (2.D) the responsiveness to changes in density. In this paper, we compare our manipulations to the null expression of the Δ*lasI*Δ*rhlI* (Figure 4) in order to test for a flattening, and therefore decreased reactivity to density, of the slope. From Figure 5 it is evident that the ‘flattening up’ hypothesis fails across all supplementation conditions, including high doses of one or both signals. Specifically, we find that in contrast to the Figure 2.C hypothesis, all supplemented reaction norm slopes are significantly greater that of the Δ*lasI*Δ*rhlI* (Dunnett’s Test, df = (15, 391) p < 0.05). We next test the ‘amplified response’ hypothesis by comparing slopes to the non-supplemented WT control and find that only the combined 5uM 3-oxo-C12 and C4 produces a significant increase in slope compared to the WT (Dunnett’s Test, df = (15, 391), one tail p = 0.0023).

Overall, from Figure 5 we can see that in most cases of signal supplementation to the WT we find that the addition of either signal alone or together (at any concentration) increases mean expression (indicated in figure by ^) while retaining a positive slope (consistent with the “robust reactivity” hypothesis, Figure 2.D). In contrast, we find no support for the ‘flatten up’ hypothesis (Figure 2.B) and minimal support for the ‘amplify’ hypothesis (indicated by *; Figure 2.C).

### Cognate and dual AHL signal complementation restores QS reactivity to density

In the next round of supplementation experiments, we now combine signal synthase knockouts with specific signal complementation (Figure 6) allowing complete experimental control over the level of one or both AHL signals experienced by cells, decoupled from density. In this context, we can now test each step of the knockout and complementation hypotheses outlined in Figure 2. Consistent with the grey lines in Figure 2.B-D, we can see that the knockouts alone lead to a relative ‘flattening off’ of the reaction norms – lower mean expression and lower slope (Figure 6 top row in grey; see Figure 4 and associated text for details on statistics).

**Figure 6.**
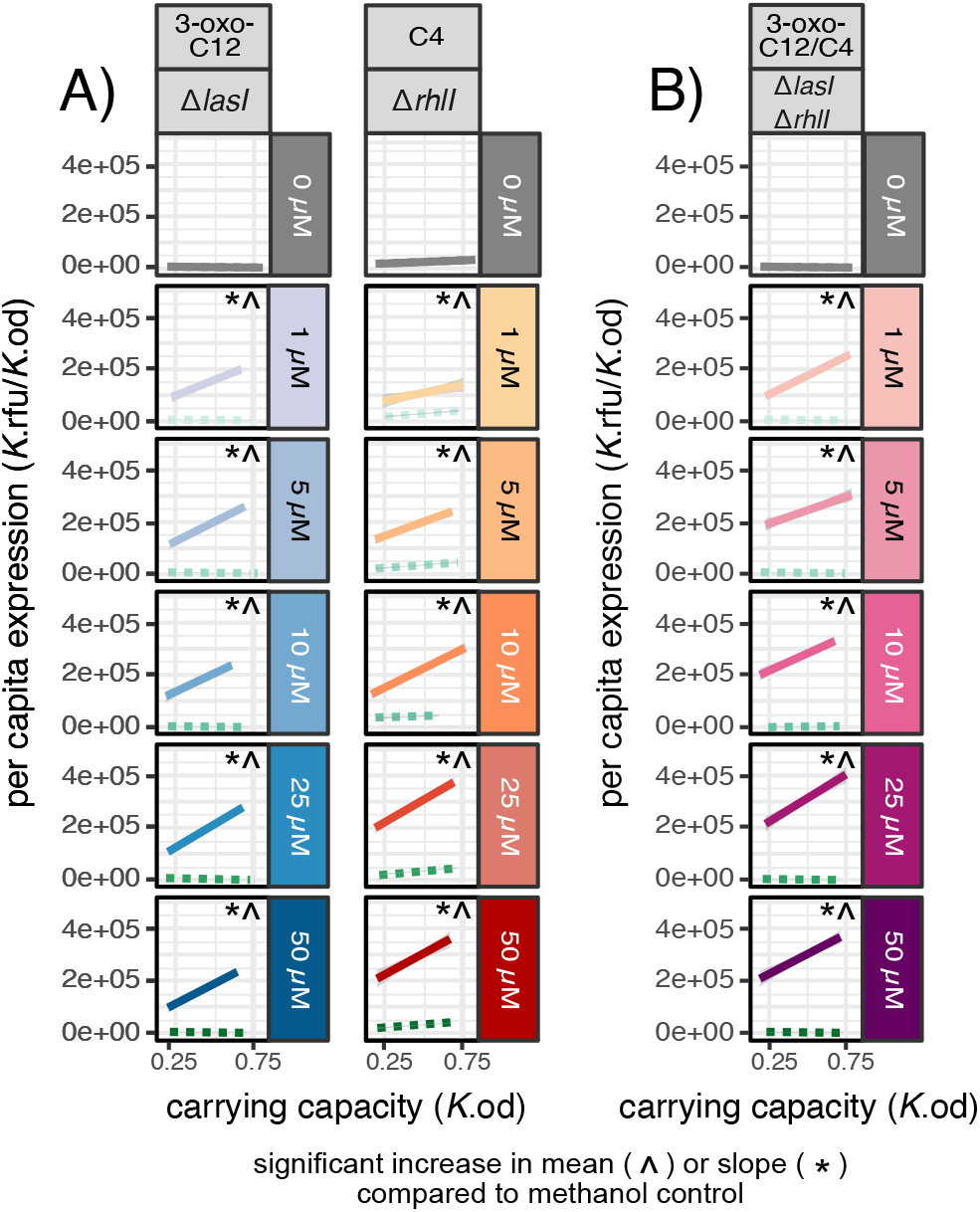
Cognate and dual AHL signal complementation restores QS reactivity to density. A) Cognate signal complementation. B) Dual signal complementation. Behavior with no signal supplementation in grey. Methanol control (green, dotted line) and signal supplemented environments [3-oxo-C12 alone (blue), C4 alone (orange), 3-oxo-C12 and C4 combined (pink)] across 5 different concentrations of signal (1uM, 5uM, 10uM, 25uM, 50uM). Each reaction norm is built using a linear regression on 18 data points (6 carrying capacity environments done in triplicate). ^ Indicates a significant increase in mean compared to the methanol control for that specific strain (Dunnett’s Test, adjusted p < 0.05) * Indicates a significant increase in slope compared to the methanol control for that specific strain (Dunnett’s Test, adjusted p < 0.05). For legibility, Figure 6 only shows the fitted linear models and not the underlying 18 datapoints per linear fit. See Supplemental Figures 1–2 for a series of plots of the experimental and control data plus linear model fits.

Our first prediction is that specifically complementing a single synthase knockout (*lasI* or *rhlI*) with cognate signal (3-oxo-C12 or C4) will restore both *lasB* expression and a positive reaction norm, due to density information being encoded by the remaining intact AHL signal synthase (Figure 2.D). Consistent with this prediction, in Figure 6.A we see that cognate signal complementation of *lasI* and *rhlI* restores QS response and a positive reaction norm. Specifically, we see that cognate signal complementation restores QS response in *lasI* (Figure 6.A, *lasI*+C12) by significantly increasing mean expression (Dunnett’s Test, df = (5,297), one tail p < 0.0001) and slope (Dunnett’s Test, df = (5,297), one tail p < 0.0001) compared to the *lasI* methanol-supplemented control (dotted green lines in Figure 6). We see similar results for cognate signal supplementation in *rhlI* (Figure 6.A, *rhlI*+C4), where QS response is also restored through a significant increase in mean (Dunnett’s Test, df = (5,274), one tail p < 0.0001) and slope (Dunnett’s Test, df = (5,274), one tail p < 0.0001) for all cases except 1uM C4 where p = 0.0168). In these cognate supplement experiments, we again can clearly see that the slope does not “flatten-up”-the slope is significantly higher than the Δ*lasI*Δ*rhlI* in Figure 4 (Dunnett’s Test, df = (5, 297), one tail, p < 0.05). This is anticipated as each strain still has a functional copy of one signal synthase gene that encodes density-*dependent* information. In terms of our “amplified response” hypothesis, while cognate signal supplementation can restore a positive slope compared to the nonsupplemented knockout, this slope is always lower than the non-supplemented WT (Dunnett’s Test, df = (10,353), one tail p < 0.001) and therefore not amplified compared to the WT (Supplemental Figure 3).

Turning to the double synthase knockout (Figure 6.B), we expect we can restore *lasB* expression via dual signal complementation, but we do not expect restoration of *lasB reactivity*, as there is no longer any connection between purely exogenously derived AHL signal supply and bacterial density. While we do see the predicted restoration of mean expression via dual signal supplementation (Dunnett’s Test, df = (5, 200), one tail p < 0.0001 for all comparisons), in contrast to the ‘flatten up’ prediction (Figure 2.B) we see that this restored expression retains a positive slope on density compared to the Δ*lasI*Δ*rhlI* (Dunnett’s Test, df = (5,200), one tail, p < 0.05). Thus, in contrast to the Figure 2.B hypothesis, we see that density-dependent *lasB* expression (a positive reaction norm) can be restored by adding a density-*independent* amount of dual signal.

### Triple signal supplementation decreases but does not ablate reactivity to density

Given that dual signal supplementation with 3-oxo-C12 and C4 does not support the ‘flatten up’ prediction, we expanded our search to other signals that could be carrying density-encoding information. The non-AHL signal molecule 2-heptyl-3-hydroxy-4(1H)-quinolone (‘<u>P</u>seudomonas <u>q</u>uinolone <u>s</u>ignal’, henceforth referred to as PQS) (50–54) is frequently linked to iron limitation and iron acquisition (51), but was first discovered as a QS signal in connection to its impact on the expression of *lasB* (50). Knocking out PQS production is more complicated than knocking out AHL production, due to the more complex genetics and pleiotropic effects of the PQS system. Choices for a genetic deletion either lead to ablating production of all 40 quinolones (Δ*pqsA*) or cell lysis through excess HHQ accumulation (Δ*pqsH*) (52). Given these choices have substantial impacts on cell behavior and fitness outside of cell-cell signaling, we manipulate PQS solely via supplementation experiments to test if PQS contains residual density-encoding information.

First, we supplement the WT with just PQS (Figure 7.A) and find that, similar to the AHLs, density-independent concentrations of PQS significantly increases the mean level of expression across the reaction norm compared to the WT (ANOVA, F(5,78) = 66, p < 2.2e-16, Dunnett’s Test, df = (5, 78), one tail p <.0001) and that in contrast to the Figure 2.B ‘flatten up’ hypothesis, all supplemented reaction norm slopes are significantly greater than the Δ*lasI*Δ*rhlI* (Dunnett’s Test, df = (5,78), one tail p < 0.05). This lack of a ‘flatten-up’ is likely due to intact production of both AHL signals (3-oxo-C12 and C4).

**Figure 7.**
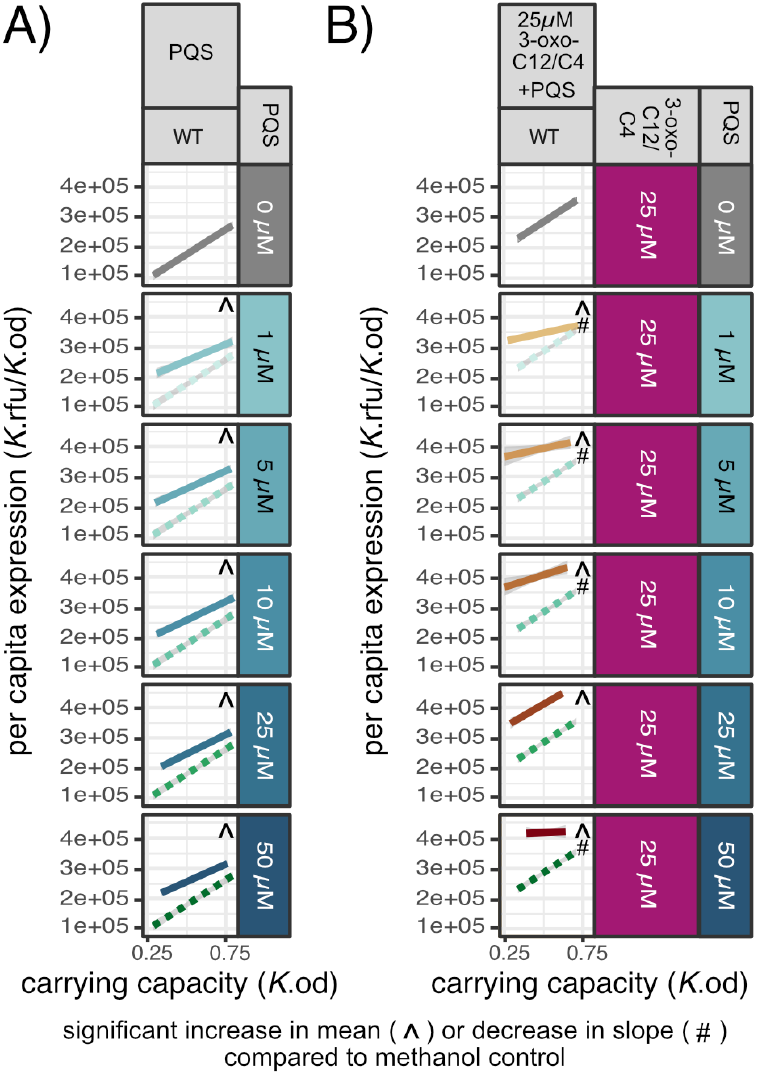
PQS and triple signal supplementation decrease, but do not always ablate, responsiveness to density. A) Variable PQS supplementation of the WT. Grey solid line represents no supplementation; green dotted lines, methanol control; teal lines, PQS signal supplemented environments. B) Fixed supplementation of 3-oxo-C12 and C4 at 25uM and variable supplementation of PQS of the WT. Grey solid line, dual 25uM AHL 3-oxo-C12 and C4 supplementation; green dotted lines, 25uM 3-oxo-C12 and C4 and methanol; brown lines, fixed 25uM 3-oxo-C12 and C4 supplementation (purple box) and variable PQS supplementation (teal box). Each reaction norm is built using a linear regression on 18 data points (6 carrying capacity environments done in triplicate). ^ Indicates a significant increase in mean compared to the methanol control for that specific strain (Dunnett’s Test, adjusted p < 0.05) # Indicates a significant *decrease* in slope compared to the methanol control for that specific strain (Dunnett’s Test, adjusted p < 0.01).

To give the best chance at a ‘flatten-up’ response, we turn to triple signal supplementation in Figure 7.B and supplement with a fixed concentration of both AHL signals and variable supplementation of PQS. We chose to supplement with 25uM of the AHLs instead of 50uM as 50uM of 3-oxo-C12 actually lowered expression across the reaction norm in Figure 5. In agreeance with Figure 5, we find that adding 25uM of 3-oxo-C12 and C4 increases expression (TEST) but not reactivity (TEST) compared to the non-supplemented WT (Figure 7.A 0uM compared to Figure 7.B 0uM).

When supplementing with all three signals, we find that supplemental PQS significantly increases the mean level of expression across the reaction norm compared to the dual AHL supplemented WT (ANOVA, F(5,72) = 106, p < 2.2e-16, Dunnett’s Test, df = (5, 72), one tail p <.0001). The increase in mean expression in the dual AHL supplemented WT upon PQS supplementation indicates that dual AHL signal supplementation alone is not sufficient to maximize *lasB* expression. But does triple supplementation of density-independent amounts of all three signals modulate the *responsiveness* of the WT to different densities? Across these triple supplementation environments, we find that only the addition of 25uM of 3-oxo-C12, 25uM of C4, and 50uM of PQS is sufficient to ‘flatten’ the reaction norm (no significant difference in slope compared to the QS null Δ*lasI*Δ*rhlI*; Dunnet’s Test, df = (5,72), p = 1). Additionally, we find no case where triple signal supplementation increases reactivity to density (slope does not increase, Dunnett’s Test, df = (5, 72), one tail p > 0.9). The results from the triple supplementation imply that only excess quantities of all three signals is sufficient to increase expression past the WT level and ‘flatten up’ the reaction norm.

## Discussion

In this study we use a reaction norm approach (Figure 2) to assess how signal knockout and supplementation treatments impact the mean QS response and *reactivity* of QS phenotypes to environmental change. Our approach condenses data from 2,000 timeseries (over 74,000 individual observations) into a comprehensive view of QS-controlled gene expression across variation in genetic, environmental, and signal determinants of *lasB* expression. Consistent with canonical understanding of QS regulatory wiring (Figure 1), we find that knocking out signal synthase genes alone and in combination attenuate the responsiveness of *lasB* to density (Figure 4). Overall, we find that both AHL and PQS signal supplementation increases expression past WT levels, indicating that native signal production is not sufficient to maximize gene expression (Figueres 4–7). We find that wildtype reactivity to density is robust to AHL and PQS supplementation (Figures 5 & 7.A) except for the most extreme case of triple supplementation (Figure 7.B). While AHL supplementation of the WT can increase overall response, it neither enhances reactivity to density nor flattens the response respective to density (Figure 5). In complementation experiments (Figure 6), we find that a positive *lasB* reaction norm is dependent on the presence of 3-oxo-C12 and can persist given density-independent AHL supplementation even in the absence of both AHL synthase genes.

A key result across our manipulations is the robustness of QS-mediated density sensing in the face of both genetic and/or chemical manipulations of signal availability (Figures 5–7). In Figure 5 we outline the robustness of the wildtype in the face of increasing exogenous supplementation with one or both AHL signals. Given we are supplementing the wildtype strain, it remains the case that at higher densities there will likely be larger concentrations of signal, due to greater contributions from the larger density of wildtype cells. While in principle this could explain the robust reactivity (consistent positive slope), we note that the preservation of a positive slope continues given supplementation with concentrations of AHL that greatly exceed observed wildtype levels (48). To remove the effect of native signal production on signal density we turned to supplementation of signal synthase knock-out strains (Figure 6), and again see robust density sensing (positive slope) phenotypes under many manipulations.

In Figure 6.B we see that a positive *lasB* reaction norm is robust even in conditions where both AHL signal concentrations are controlled independently of bacterial density. This points to a role for additional density-dependent factors controlling *lasB* expression, outside of 3-oxo-C12 and C4 HSL signal density. Figure 7.B indicates that, in the extreme case, supplementing with PQS can ablate the ability to sense density. Recent work points to PqsE, an effector protein of PA QS, as an additional control dial for *lasB* (53–55). While PqsE is not required for RhlR-driven gene expression, Letizia et al. 2022 show that PqsE can modulate the level of expression of RhlR controlled genes, such as *lasB*. In addition to PQS, *lasB* expression is impacted by multiple other factors, which in turn may be impacted by other aspects of density manipulations. For example, by manipulating the availability of limiting carbon (Figure 3.C), we also modify the time and number of generations until arrival at stationary phase.

From an evolutionary perspective, we can view our manipulative experiments in the context of animal communication theory (57,58). Communication systems convey reliable information when signals correlate with underlying information of interest (59). For example, the jumping height of ‘stotting’ gazelles signals to predators reliable information on the athletic ability of potential prey, leading predators to avoid pursuit of the most athletic (high-stotting) individuals (60). One important challenge to signal reliability is noise – environmental forces that degrade or distort the transmitted signal and therefore weaken the correlation with useful information (e.g. factors reducing visibility in the context of stotting signals). Evolutionary theory predicts that if signal reliability is reduced by noise, communication breaks down as receivers are selected to ignore the signal and signalers respond by dropping signal production (59,61,62). Partially consistent with this basic prediction, Popat et al. 2015 showed that long-term QS signal supplementation (50μm of 3-oxo-C12; described as adding uncorrelated noise to QS communication) over 120 generations in *P. aeruginosa* selected for an evolved response of reduced signal investment. However, Popat et al. 2015 did not examine behavior across reaction norms so we do not know to what extent the signal manipulation functioned as ‘noise’, i.e. to what extent it obscured the relationship between density, QS signal and QS response. Our experimental results shed light on this issue and highlight that this supplementation design likely elevated average response, while maintaining responsiveness of the ancestral *P. aeruginosa* to changes in density, potentially leading to early and excessive production of *lasB* (required in Popat et al.’s experiment in order to digest the carbon source). Our findings in Figure 5 indicate that *P. aeruginosa* QS populations similar to our strain may actually be fairly robust to excess exogenous signal and are still able to sense density when challenged with density-*independent* signal.

One limitation of our experimental method is that quantifying QS activity from batch culture does not represent true steady state dynamics. Chemostats present an alternative approach that offer a more controlled steady state environment, although our ability to assess multiple distinct treatments would then be limited by the larger spatial scale and complexity of chemostat approaches. While we cover a 20-fold range of density in this manuscript (the range of cell densities generated from our culture method is roughly 1×10^8^ cells/mL to 2x 10^9^ cells/mL), chemostats, or at least larger batch cultures than microtiter plates, would also increase the range of densities that are observable. In an effort to make our results translatable across strains of PAO1, we quantified both AHLs in our highest density environment and find that the levels of signal (0.8uM 3-oxo-C12 and 2.3uM C4) agree with other high density work done in the field. In addition to signal, it would also be interesting to look at what other factors impact the ability of the WT to sense density even when given a density-*independent* concentration of signal. This could be done using constructs containing inducible transcription factors like *lasR* to increase *lasR* expression, or by using *rhlR* specific C4 competitive binders to reduce *rhlR* mediated activity. Transcriptomics could provide a good baseline as a discovery step to determine which mechanistic avenues would be promising to pursue via controlled reaction-norm experiments.

Overall, our results show that a positive reaction norm to density is robust to multiple combinations of gene deletion and density-independent signal supplementation. Our work develops a modular approach to query the robustness and mechanistic bases of the central environmental *sensing* phenotype of quorum sensing.

## Methods

### Bacterial Strains and Growth Conditions

The main bacterial strain used in this study is *P. aeruginosa* NPAO1 (Nottingham-PAO1) containing the *PlasB::gfp(ASV*) quorum sensing reporter pMHLAS (64). Single and double signal synthase knockouts (Δ*lasI*, Δ*rhlI*, Δ*lasI*Δ*rhlI*) were made using double allelic exchange and the quorum sensing reporter pMHLAS was subsequently electroporated in. Overnight cultures were grown in lysogeny broth (LB), supplemented with 30 ug/ml gentamicin to maintain the pMHLAS plasmid, with shaking at 37 °C. Experiments were conducted in lightly buffered (50 mM MOPS) M9 minimal defined media composed of an autoclaved basal salts solution (Na_2_HPO_4_, 6.8 gL^−1^; KH_2_PO_4_, 3.0 gL^−1^; NaCl, 0.5 gL^−1^), and filter-sterilized 1 mM MgSO_4_, 100 uM CaCl_2_, and 1X Hutner’s Trace Elements with casein digest, as the sole carbon source (ThermoFisher Difco™ Casein Digest CAT 211610).

### Controlling Culture Carrying Capacity

We manipulated density by controlling the limiting resource in the media, carbon, allowing us to tune the carrying capacity of each treatment and verified that carbon was the limiting resource across our range of densities. To cover a variety of densities, we generated a carbon range between 0.05% and 0.25% via dilutions of a 0.5% carbon minimal media stock for a total of 6 different carrying capacities. Quantities of carbon past 0.25% start exhibiting characteristics of non-logistic growth, so we use 0.25% to generate our highest densities. This produced a range of densities environments from 1.18×10^8^ cells/ml to 2.02×10^9^ cells/ml. Overnight cultures were grown in LB gentamicin 50 ug/ml and centrifuged for 2 minutes. The cells were then washed twice with carbonless minimal media and then each carbon treatment was adjusted to OD600 = 0.05. Then, 200 uL of each sample was added to a 96-well microplate with the buffered M9 minimal media described above. Plates were incubated with shaking at 37 °C in a Cytation/BioSpa™ plate reader with readings of optical density (OD_600_) and green fluorescence taken at 30-min intervals for 20 hours.

### Estimating per capita lasB expression

To estimate per capita *lasB* expression, we first derive steady-state estimates of population density (*K_OD_*) and and population fluorescence (*K_RFU_*), by fitting logistic curves (growthcurver() package in R) to the OD and RFU timeseries data over 20 hours (see Figure 3A). We then estimate per-capita *lasB* expression as the ratio of these values, *K_RFU_/K_OD_*. To produce reaction norms, we plot per capita expression (*K_RFU_/K_OD_*) against carrying capacity (*K_OD_*) (Figures 4–7).

### Exogenous signal addition

The AHL signaling molecules used in this study are N-(3-oxododecanoyl)-L-homoserine lactone (3-oxo-C12-HSL, Sigma-Aldrich CAS# 168982-69-2), N-butyryl-homoserine lactone (C4-HSL, Sigma-Aldrich CAS# 98426-48-3), and 2-Heptyl-3-hydroxy-4(1H)-quinolone (pqs, Sigma-Aldrich CAS# 108985-27-9) alone and in combination. These signals are suspended in methanol, so methanol was used as a control to account for different amounts of signal being added experimentally (Supplemental Figure 2).

### Statistical Analysis

Statistical analysis was performed using R. Logistic curves were fit to raw expression (RFU) and growth (OD_600_) data using the growthcurver package. *K*, commonly called carrying capacity, was extracted from the logistic model. Linear models were then built from the extracted *K* values using the emmeans package. Both means and slopes were obtained from those linear models using the emmeans() function and emtrends() function which uses estimated marginal means to construct a reference grid of predicted means or trends. ANOVA was used for pairwise comparisons between all treatments. Dunnett’s Test for Multiple Comparisons was used to compare the manipulations with their respective controls (defined in their respective results sections). Reported p values are adjusted for multiple comparisons using the Šídák Method. Statistical tests for hypotheses with specific directionality (i.e., a statistically significant increase or decrease) are denoted in the text as one tailed.

## Supplemental Materials

**Supplemental Figure 1.**
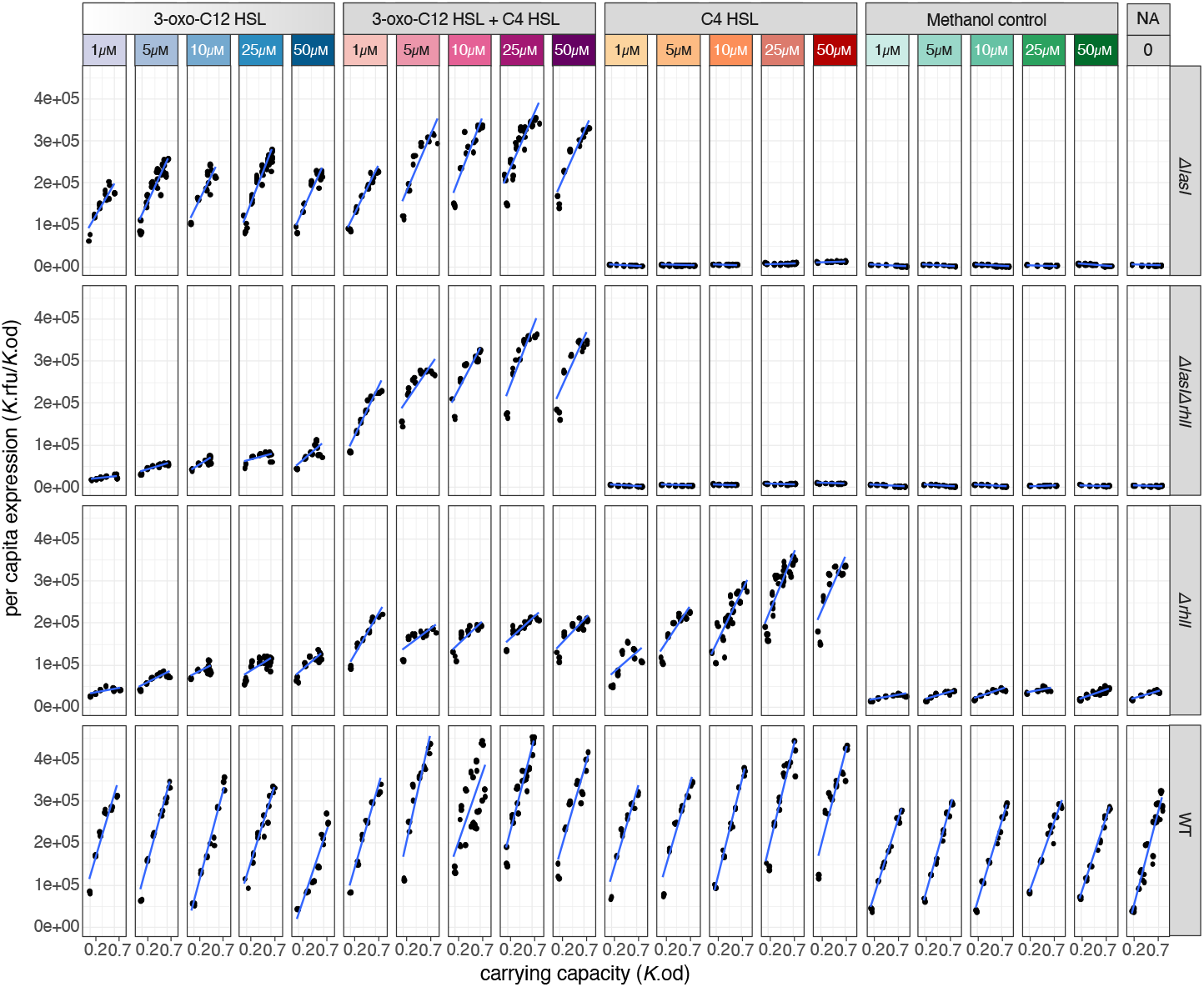
Data and linear model fits. Data (black circles) with linear model fits (blue lines). Each dot summarizes a 20-hour timeseries experiment (see Figure 2).

**Supplemental Figure 2.**
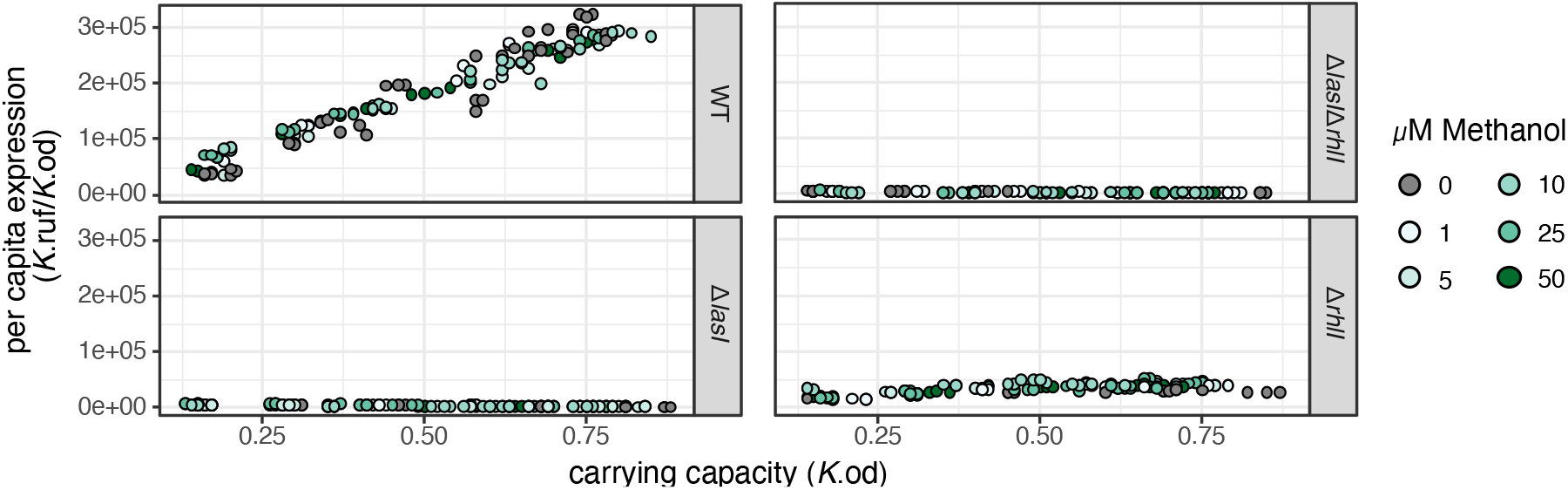
Methanol does not impact expression or growth. In the case of supplementing signal, the AHL signaling molecules used in this study are suspended in methanol. Therefore, we first tested if different concentrations of methanol impacted our experiments and found that overall, methanol did not significantly impact (ANOVA, df = 500, p = 0.627) or expression (ANOVA, df = 500, p = 0.133).

**Supplemental Figure 3.**
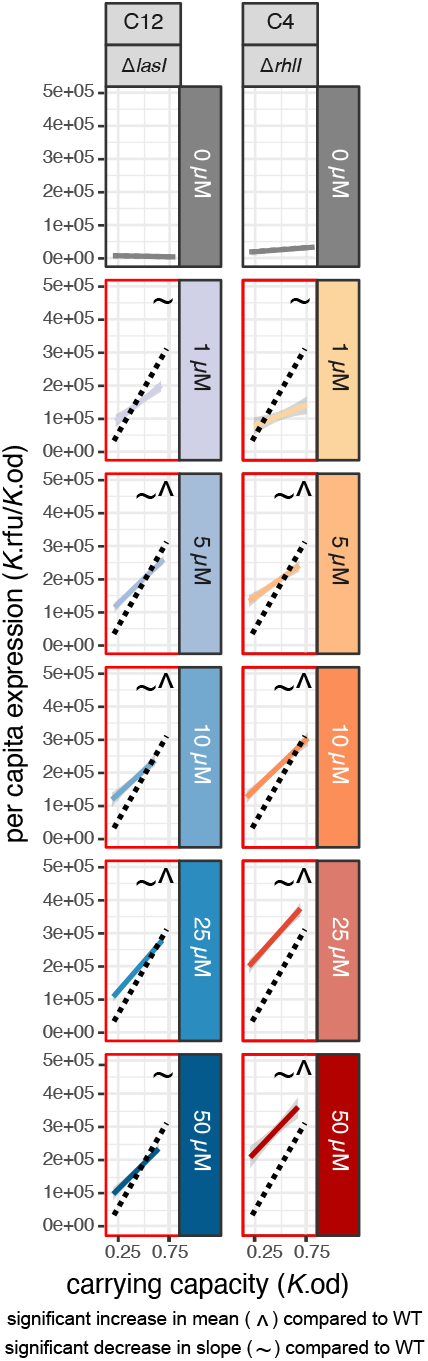
Cognate signal supplementation compared to the WT expression. In most comparisons, mean expression is significantly higher than WT (Dunnett’s Test, df = (10,353), p_lasI+50uM3-oxo-C12_ = 0.543, p_lasI+1uM3-oxo-C12_ = 1, P_rhlI+1uMC4_ = 1, otherwise p < 0.001), but slope is significantly lower compared to the WT (Dunnett’s Test, df = (10,353), p < 0.001 for all comparisons).

